# *In Vivo* Mechanisms of Chemotherapy-Induced Acute Follicle Loss in the Human Ovary: An Individual-Oocyte Transcriptomic Analysis from Human Ovarian Xenografts

**DOI:** 10.1101/2020.06.23.167783

**Authors:** S. Titus, K.J. Szymanska, B. Musul, V. Turan, E. Taylan, R. Garcia-Milian, S. Mehta, K. Oktay

## Abstract

Gonadotoxic chemotherapeutics, such as cyclophosphamide, cause early menopause and infertility in women. Earlier histological studies showed ovarian reserve depletion via severe DNA damage and apoptosis, but others suggested activation of PI3K/PTEN/Akt pathway and follicle ‘burn-out’ as a cause. Using a human ovarian xenograft model, we performed single-cell RNA-sequencing on laser-captured individual primordial follicle oocytes 12h after a single cyclophosphamide injection to determine the mechanisms of acute follicle loss after gonadotoxic chemotherapy. RNA-sequencing showed 190 differentially expressed genes between the cyclophosphamide- and vehicle-exposed oocytes. Ingenuity Pathway Analysis predicted a significant decrease in the expression of anti-apoptotic pro-Akt *PECAM1* (p=2.13E-09), *IKBKE* (p=0.0001), and *ANGPT1* (p=0.003), and reduced activation of PI3K/PTEN/Akt after cyclophosphamide. The qRT-PCR and immunostaining confirmed that in primordial follicle oocytes, cyclophosphamide did not change the expressions of *Akt* (p=0.9), *rpS6* (p=0.3), *Foxo3a* (p=0.12) and anti-apoptotic *Bcl2* (p=0.17), nor affect their phosphorylation status. There was significantly increased DNA damage by γH2AX (p=0.0002) and apoptosis by active-caspase-3 (p=0.0001) staining in the primordial follicles and no change in the growing follicles 12h after chemotherapy. These data suggest that the mechanism of acute follicle loss by cyclophosphamide is via apoptosis, rather than growth activation of primordial follicle oocytes in the human ovary.

**One Sentence Summary:** Single-cell transcriptomic interrogation of primordial follicles in human ovarian xenografts reveals that chemotherapy causes acute ovarian reserve depletion by inducing a pro-apoptotic state rather than activating pathways that result in follicle growth initiation.

## INTRODUCTION

In 2019, approximately 891,480 women were diagnosed with cancer in the U.S. alone, of whom 5.5% were in reproductive age [1], and potentially received fertility-damaging cancer treatments. Worldwide, chemotherapy-induced premature menopause and infertility are significant quality of life issues for young women diagnosed with cancer. To preserve fertility in the face of gonadotoxic chemotherapy, successful gamete, and gonadal tissue cryopreservation techniques have been developed [2]. However, these require at least minimally invasive procedures, and there have been no proven targeted medical treatments to block the ovarian damaging side effects of chemotherapy. If the mechanisms of chemotherapy-induced ovarian damage are understood, noninvasive, targeted pharmacological methods of fertility preservation can be developed.

Based on our earlier work, we suggest that chemotherapy causes both acute and delayed effects on the primordial follicle reserve. Chemotherapy agents such as doxorubicin, a topoisomerase inhibitor [3], and cyclophosphamide, an alkylating agent [4][5], cause apoptotic primordial follicle death by inducing DNA double-strand breaks (DSBs) in the human ovary. We found that the DNA damage peaks at around 12h and subsequent follicle loss at 48h, accounting for the acute loss of follicle reserve. Besides, we also showed in *in vivo* human ovarian xenograft models that doxorubicin causes stromal cell death as well as microvascular damage, inducing tissue hypoxia, which may contribute to the delayed loss of ovarian follicles. Others have also shown possible stromal damaging effects of cancer drugs in women [6]. It has also been proposed in rodent models that cyclophosphamide may induce depletion of primordial follicle reserve via inducing a massive entry of follicles into the growth phase [7]. It was surmised that this follicle ‘burn-out’ is the acute result of the direct activation of phosphoinositide 3-kinase/phosphatase and tensin homolog/protein kinase B (PI3K/PTEN/Akt) pathways, which regulate primordial follicle growth initiation. There could be several reasons for these discrepancies, including the possibility of the differing mechanism being in play at acute and delayed phases after the chemotherapy exposure [8]. Therefore, mechanistic studies are needed to delineate the acute impact of chemotherapy on ovarian follicle reserve. Utilizing the human ovarian xenografting coupled with laser-capture-microdissection-assisted individual-oocyte RNA-sequencing (RNA-seq) and quantitative real-time PCR (qRT-PCR) approaches perfected by our team, we performed this study to determine the molecular mechanisms of acute chemotherapy-induced loss of primordial follicles in the human ovary [4][5].

## MATERIALS AND METHODS

### Mice

Female severe combined immunodeficiency (NOD-SCID) mice aged 2-3 months were obtained from Taconic farms and housed under pathogen-free conditions. Animals were kept in a 12 h light-dark cycle and provided ad libitum with water and a laboratory diet. All animal experiments were approved and conducted in accordance with the protocol of the Yale University Institutional Animal Care and Use Committee.

### Ovarian Xenografts

Ovarian cortical pieces from cadaveric organ donors aged ≤ 32 years were cryopreserved according to our established protocol [9,10]. Tissue pieces 2-3 mm^3^ (2-3 × 1 × 1 mm) were xenografted subcutaneously to SCID mice, as described previously [4]. Briefly, after animals were anesthetized, a small incision was made on the dorsal midline of the animal. Human ovarian cortex fragments were transplanted to each animal (n=4/animal). After allowing xenografts to fully vascularize for 10 days, the mice were given a single intraperitoneal (i.p.) dose of cyclophosphamide (75 mg/kg) or the vehicle. The tissues were then recovered 12h later and prepared for either histology and immunohistochemistry or embedded in a medium for frozen sectioning and isolation of primordial follicle oocytes by laser capture dissection microscopy.

### Immunohistochemistry

Ovarian tissues were fixed in 4% paraformaldehyde (PFA), embedded in paraffin, and serially sectioned at 5-micron intervals. Sections were stained with Hematoxylin and Eosin (H&E) for differential follicle counts, as previously described [4]. Every fifth section of an entire xenograft was chosen for staining and evaluated by observers who were blinded to the intervention and each other’s findings. To avoid over-counting, follicles were recorded only when the nucleus was visible. Follicles were classified based on our previously established criteria [11]. A primordial follicle was defined as an oocyte surrounded by a single layer of flattened, squamous follicular cells. A primary follicle was defined as an oocyte surrounded by a single layer of cuboidal granulosa cells. A follicle was classified atretic based on morphological criteria, including nuclear pyknosis, cytoplasm contraction, and nuclear pyknosis of granulosa cells and dissociation of the granulosa cells from the basal membrane [12,13].

Randomly selected sections coming from ovarian tissue of least 3 different individuals were used for immunohistochemistry for Anti-active Caspase-3 (AC-3, 1:1000, AF-835, R&D Systems), γH2AX (1:250 IHC-00059, Bethyl Laboratories), rabbit phospho-Akt (S473, 1:400, #4060S, Cell Signaling Technology, Denver, MA), rabbit phospho-rpS6 (S235-236, 1:100; #2211S, Cell Signaling Technology, Denver, MA), and phospho-FOXO3A (S253, 1µg/ml, # ab47285, Abcam, Cambridge, MA, USA). Images were captured on an Olympus IX73 microscope. BAD (1:250, A302-384A, Bethyl Laboratories) and Bcl2 (1:200, sc-509, Santa Cruz Biotechnology). Nikon 90i Eclipse microscope was used for imaging. By overlapping both BAD and Bcl2 staining color channels, and selecting the area around primordial follicles, we calculated the mean pixel intensities. We compared the intensity means between primordial follicle oocytes of vehicle and chemo groups after subtracting the background by setting a constant threshold for both channels across the images. For the calculation, we used 3 slides of each xenografted tissue, and ‘n’ represents the number of different tissue pieces. The analysis was done using ImageJ software.

### Laser-capture microdissection (LCM) and isolation of single primordial follicle oocytes

After harvesting ovarian xenografts, tissue blocks were immediately prepared in optimum cutting temperature (OCT) compound and stored at −80°C until cryostat sectioning (−20°C). These were later sectioned at 8-micron thickness and stained with H&E according to the manufacturer’s protocol. Based on the morphological appearance, i.e., follicle with a single layer of flat granulosa cells around with positively stained oocyte inside, single primordial follicle oocytes were individually isolated using an LCM excluding any pre-granulosa cell from the dissection. To each Eppendorf tube, filled with cell lysis buffer (RNase inhibitor in 0.2% (vol/vol) Triton X-100 solution), the individual primordial oocyte was collected from 2-3 adjacent tissue sections to ensure that one full cell was included (Leica Microsystems, Buffalo Grove, IL, USA).

### RNA-seq analysis on laser captured single primordial follicle oocytes

The primordial follicle oocytes were processed for sequencing with modifications of a previously established protocol on non-germline single cells for LCM material [14]. Samples were sequenced to depths of up to 44 million read pairs, 75 nt length reads per sample using the Illumina Rapid v2 kit (75 cycles) on an Illumina HiSeq2500 Sequencing System. Image analysis, base calling, and generation of sequence reads were produced using the HiSeq Control Software v2.0 (NCS) and Real-Time Analysis Software (RTA). Data were converted to FASTQ files using the bcl2fastq2 v1.8.4 software (Illumina Inc.). The reads were trimmed for quality. Those were then aligned to the human reference genome (hg38 gencode for humans) using HiSAT2 [15]. The alignments were processed using Ballgown (free online software) [15], and per-gene counts were obtained. The raw counts were processed using DESeq2 and R.

### Gene network and pathway analysis

Ingenuity Pathway Analysis (IPA) Ingenuity Systems QIAGEN, Content version: 45865156, 2018, Redwood City, CA, USA) was used to carry out pathway analysis for differentially expressed genes across samples. Each gene symbol was mapped to its corresponding gene object in the Ingenuity Pathways Knowledge Base. The over-represented pathways are ranked according to the calculated *Benjamini-Hochberg* FDR P<0.05 [16].

### qRT-PCR on laser captured primordial follicle oocytes

RNA from single primordial follicles was amplified, and cDNA was prepared using established protocol [14], and gene expression analysis was done by qRT-PCR using Syber Green (Applied Biosystems Warrington, UK) on ABI QuantStudio-6 Flex Real-Time PCR machine. PCR cycling conditions were 95°C for 5 min and 45 cycles of 10 s at 95°C, 58°C for 15 s, and 72°C for 15 s, followed by a melting curve analysis to confirm the single, specific products. The expression level of each mRNA was calculated by the comparative Ct method (ΔΔCt), and the fold change (FC) was calculated by the equation 2^−ΔΔCt^. β-Actin was used as a reference gene for normalization. Primers for qRT-PCR reactions are specified in Supplementary Table 1 and were obtained from UDT (USA). ‘n’ indicates the number of single follicles. Each experiment contained primordial follicles from at least 3 different tissue donors.

### Statistics

For RNA-seq analysis, we determined statistical significance for differential gene expression based on the q-value (or the False Discovery Rate), with stringent cut-offs of 0.05 or less. This ensures that the Type II error is kept as low as possible, given the constraints of sample size. Our observations of gene transcript abundances by RNA-seq from a relatively low sample-size are justified as we observe similar trends while interrogating the samples using other more direct biological and biochemical approaches such as qRT-PCR analysis and immunohistochemistry. Other data from chemotherapy- and vehicle-treated groups were compared using Student’s two-tailed t-test. P values less than 0.05 were considered significant.

## RESULTS

### Cyclophosphamide treatment does not increase primordial follicle growth initiation; it induces DNA DSBs and apoptotic death

First, to morphologically determine if cyclophosphamide exposure increases the entry of primordial follicles into the growth pool, we calculated the ratio of primary to primordial follicles (py/pd = follicle growth initiation index) in xenografts [17]. Based on the growth initiation ratio, we are only able to give an estimate, and not a definite answer of the follicle fate since we use only a small fragment of the cortex and of the unequal distribution of the follicles in each tissue. For H&E and AC-3 staining, we analyzed a total of 70 follicles (51 primordial and 19 primaries) from 3 slides of 8 different tissue pieces for the vehicle group and 202 follicles (177 primordial and 25 primaries) from 3 slides of 10 pieces of cortex from the chemo-treated group. We found no significant difference in the py/pd ratio between the vehicle and the cyclophosphamide -treated group (34.62 ± 7.94, vs. 19.97 ± 6.79; p = 0.17) (Figure 1A). Because atretic follicles may not initiate growth, we also analyzed the ratio of non-atretic py/pd follicles. We, again, did not observe a significant difference in follicle growth initiation rate between the vehicle- and cyclophosphamide -treated groups (34.59 ± 9.75 vs. 18.58 ± 10.21; p = 0.08) (Figure 1B).

**Figure 1.**
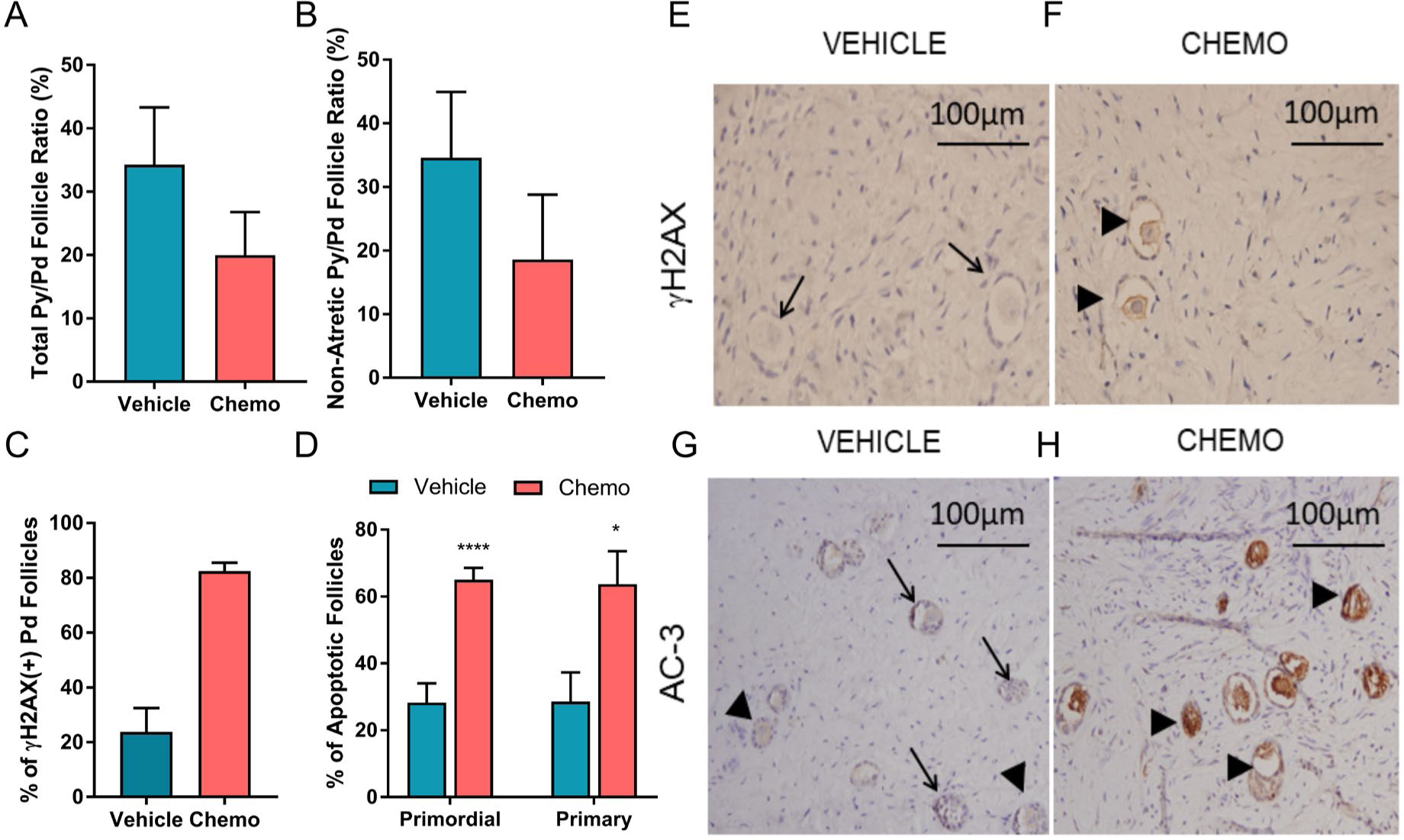
Cyclophosphamide does not trigger follicle growth initiation but induces DNA DSBs and apoptosis in human primordial follicles 12h after exposure. There was no significant difference observed in (A) total (p = 0.17), (B) non-atretic (p = 0.08) primordial follicle growth initiation rate as calculated by primary to primordial (py/pd) follicle ratios (n_vehicle_ = 8, n_chemo_ = 10). (C) Chemotherapy treatment induced significant DNA damage (γH2AX positive +) in primordial follicles (p = 0.0001). (D) There was a significant increase in the percentage of apoptotic primordial (p = 0.001) and primary (p = 0.01) follicles in cyclophosphamide -treated samples (n_vehicle_ = 8, n_chemo_ = 10). Photomicrographs of γH2AX (E-F) and AC-3 staining (G-H) positive follicles in vehicle- and cyclophosphamide -treated (chemo) groups. Black arrows indicate negative, black arrowheads show follicles positive for AC-3 and γH2AX. The ‘n’ equals the number of xenografted ovarian tissue, data are presented as mean ± S.E.M.

Next, we studied whether cyclophosphamide induces apoptotic oocyte death using AC-3 immunostaining. We found a significant increase in the percentage of apoptotic primordial (65.03% ± 3.55% vs. 28.27% ± 5.77%; p < 0.0001 cyclophosphamide vs vehicle) and primary follicles (63.67% ± 9.89%, n = 8 vs. 28.57% ± 8.74%; n = 10; p = 0.01 cyclophosphamide vs vehicle) indicating that cyclophosphamide caused massive apoptotic follicle death within 12h of treatment (Figure 1C-F). Moreover, confirming earlier reports, we found by γH2AX staining that chemotherapy induced DNA DSBs in primordial follicles compared to the vehicle treatment (82 % ± 3.1%, n = 5 vs. 23.82% ± 8.66%, n = 5; p = 0.0002).’n’ equals the number of xenografted tissue (Figure 1G-H).

### Individual-oocyte RNA-Seq Analysis of Pathways Activated in Primordial Follicles in Acute Response to Chemotherapy

We then performed high throughput RNA-seq on non-atretic, individual primordial follicle oocytes from cyclophosphamide - and vehicle-treated xenografts to have a broader view of the pathways that are altered in primordial follicles in acute response to chemotherapy. For each condition, we used two individual primordial follicles from each of the two different donor ovaries. On average, 29.8 mln reads were obtained per sample with 74% alignment to the human genome. We found that 190 genes were differentially expressed between the chemo- and vehicle-treated groups (fold change ≥ 2, p < 0.05) (Figure 2A and Supplementary Table 2). We then performed IPA enrichment analysis (Qiagen) to determine significantly altered (Figure 2B) canonical pathways (p = 0.05; fold change [fc] = 2). We found that with chemo-exposure, the Fcγ Receptor-mediated Phagocytosis in Macrophages and Monocytes (p = 0.005), Phospholipase C Signaling (p = 0.005), Ephrin Receptor Signaling (p = 0.005) and Interleukin-8 signaling (p = 0.03) were suppressed. We also found significant changes in Axonal Guidance Signaling (p = 0.0003), Gap junction signaling (p = 0.01), and Focal Adhesion Kinase (FAK) Signaling (p = 0.04), but no predictions could be made as to the direction of change. The Phospholipase C Signaling, Ephrin Receptor Signaling, and IL8 signaling pathways are known to be associated with inhibition of apoptosis. We used IPA to generate the network connecting the genes present in these pathways with the apoptosis function (p = 5.37E-19) and to predict if apoptosis is favored based on the expression status of these genes in our data. The IPA suggested activation of apoptotic processes in the chemotherapy-treated samples (Supplementary Figure1). Further, we used IPA to find the overlap between the differentially expressed genes after chemotherapy exposure and those genes in the IPA knowledge base associated with the functions of apoptosis and activation of primordial follicles. We found in our data set that *PTEN, Akt-1 Foxo3, BAD, ANGPT1, Pde* and *BTG2* overlapped with apoptotic function (p = 3.58E-18), while *PTEN* and *KITLG* overlapped with the activation function of ovarian follicles (p = 7.90E-07). The analysis of our dataset showed that the growth activation of the primordial follicle is inhibited while apoptosis is triggered in acute response to chemotherapy (Figure 2C).

**Figure 2.**
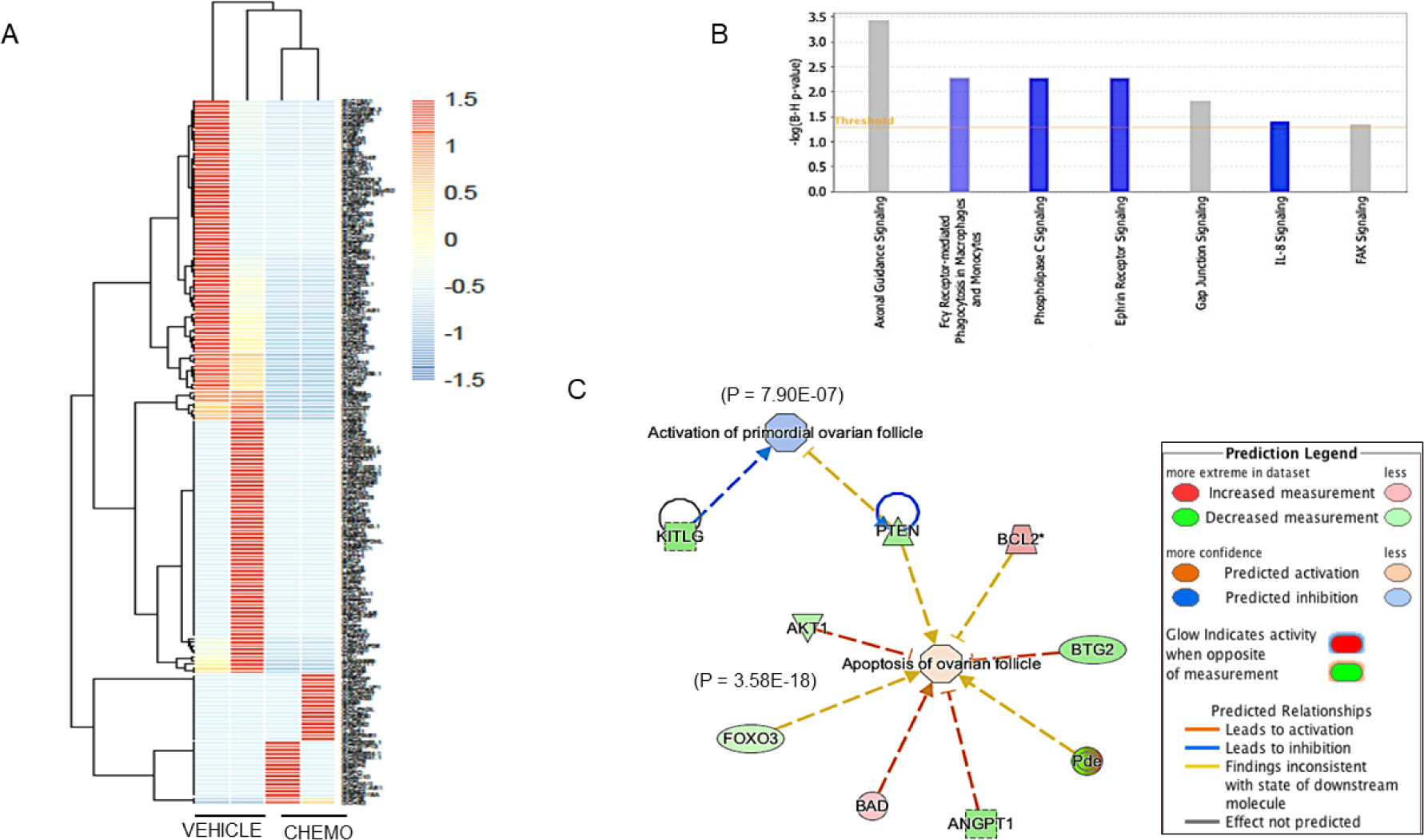
Individual-oocyte RNA-Sequencing and Ingenuity Pathway Analysis (IPA) Results. (A) Heat map of genes that are significantly altered 12h after cyclophosphamide exposure compared to the vehicle treatment. (B) Major pathways that were found to be significantly altered after chemotherapy exposure. The over-represented pathways are ranked according to the calculated *Benjamini-Hochberg* FDR, p<0.05. (C) IPA was utilized to find the overlap between the differentially expressed genes in our study and those genes in the IPA knowledge base associated with the functions of apoptosis and activation of primordial follicles. This analysis predicted relationships that favored apoptosis and inhibition of the Akt pathway in primordial follicles 12h after chemotherapy exposure.

Supporting these findings, RNA-seq analysis showed that cyclophosphamide did not cause transcriptomic activation PI3K/PTEN/Akt pathway genes, which are responsible for primordial follicle growth initiation. Specifically, RNA-seq analysis showed a significant decrease in the expressions of *PECAM1* (p = 2.13E-09), *IKBKE* (p = 0.001) and *ANGPT1* (p = 0.03) (Figure 3A-C) in the cyclophosphamide -treated samples when compared to the vehicle. These genes have anti-apoptotic functions and are also known activators of Akt [18][19][20][21][22]. Further analysis of the PI3K/PTEN/Akt pathway in the cyclophosphamide -treated samples showed a trend toward a decline in *Akt-1* (p = 0.19) and no significant change in the expressions of *Foxo3a* (p = 0.8) and *rpS6* (p = 0.36) (Figure 3D-F). These findings suggest that cyclophosphamide treatment does not result in the activation of follicle growth or anti-apoptotic pathways but triggers an acute apoptotic response.

**Figure 3.**
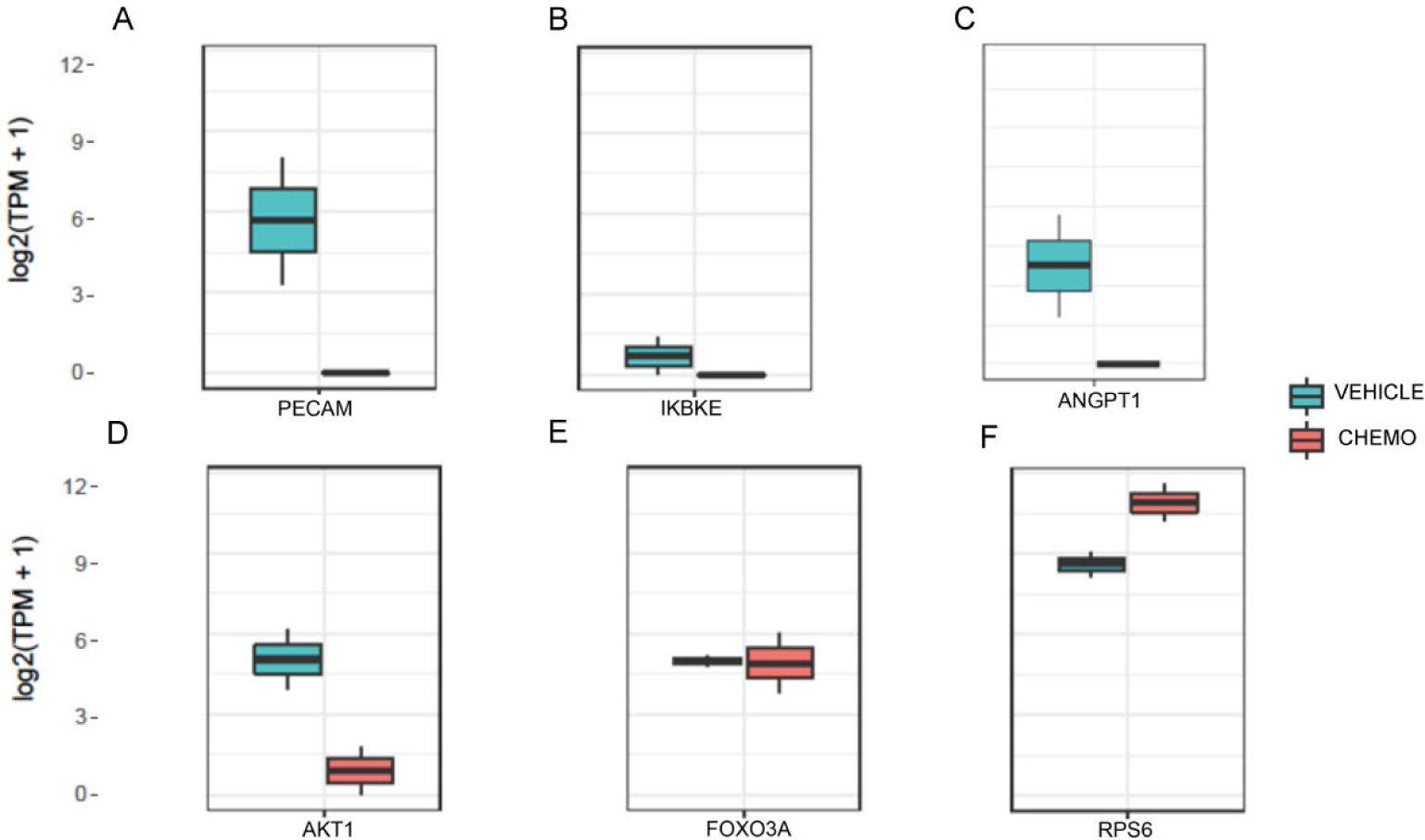
RNA-Sequencing results indicating lack of activation of the PI3K/PTEN/Akt pathway 12h after chemotherapy exposure. Significant decrease in the expression of (A) *PECAM1* (p = 2.13E-09), (B) *IKBKE* (p = 0.001) and (C) *ANGPT1* (p = 0.03) that are reported to be anti-apoptotic with pro effects on the PI3K/PTEN/Akt pathway. In cyclophosphamide - treated samples, there was a trend towards a decline in the expression of PI3K/PTEN/Akt pathway member *Akt-1* (p = 0.19) (D) while no changes in the expressions of the *Foxo3a* (p = 0.8) (E) or *rpS6* (p = 0.36) (F) were observed.

### qRT-PCR and Immunohistochemical Confirmation of Chemotherapy-Induced Acute Pathway Changes in Human Primordial Follicles

To further explore and confirm the expression patterns of the PI3K/PTEN/Akt pathway members in primordial follicle oocytes in acute response to chemotherapy, we performed qRT-PCR analysis on LCM, individual primordial follicle oocytes. Based on the relative gene expression analysis, we found no change in the expressions of *Akt-1* (n _vehicle_ = 5 and n _chemo_ = 6; p = 0.93), *rpS6* (n _vehicle_ = 5 and n _chemo_ = 6; p = 0.32) and *Foxo3a* expression (n _vehicle_ = 5 and n _chemo_ = 6; p = 0.12) 12h after cyclophosphamide exposure compared to the vehicle (Figure 4A). To further confirm our findings, we investigated the changes in the immunohistochemical expression of the key PI3K/PTEN/Akt pathway proteins in response to chemotherapy (Figure 4B). We calculated the percentage of positive to the total number of follicles from 3 slides per xenografted tissue, ‘n’ indicating the number of tissue pieces. We observed no significant differences between the vehicle and cyclophosphamide treatment groups in the expressions of p(phospho)-Akt (7.29 ± 0.22, n = 8 vs. 11.59 ± 0.8, n = 6; p = 0.52), p-Foxo3a (12.03 ± 1.48, n = 9 vs. 18.85 ± 1.43, n = 8 ; p = 0.63), and p-rpS6 (6.06 ± 0.14, n = 11 vs.12.55 ± 0.94, n = 7; p = 0.4). These results further confirm that chemotherapy does not activate the PI3K/PTEN/Akt pathway in primordial follicles within 12h of exposure.

**Figure 4.**
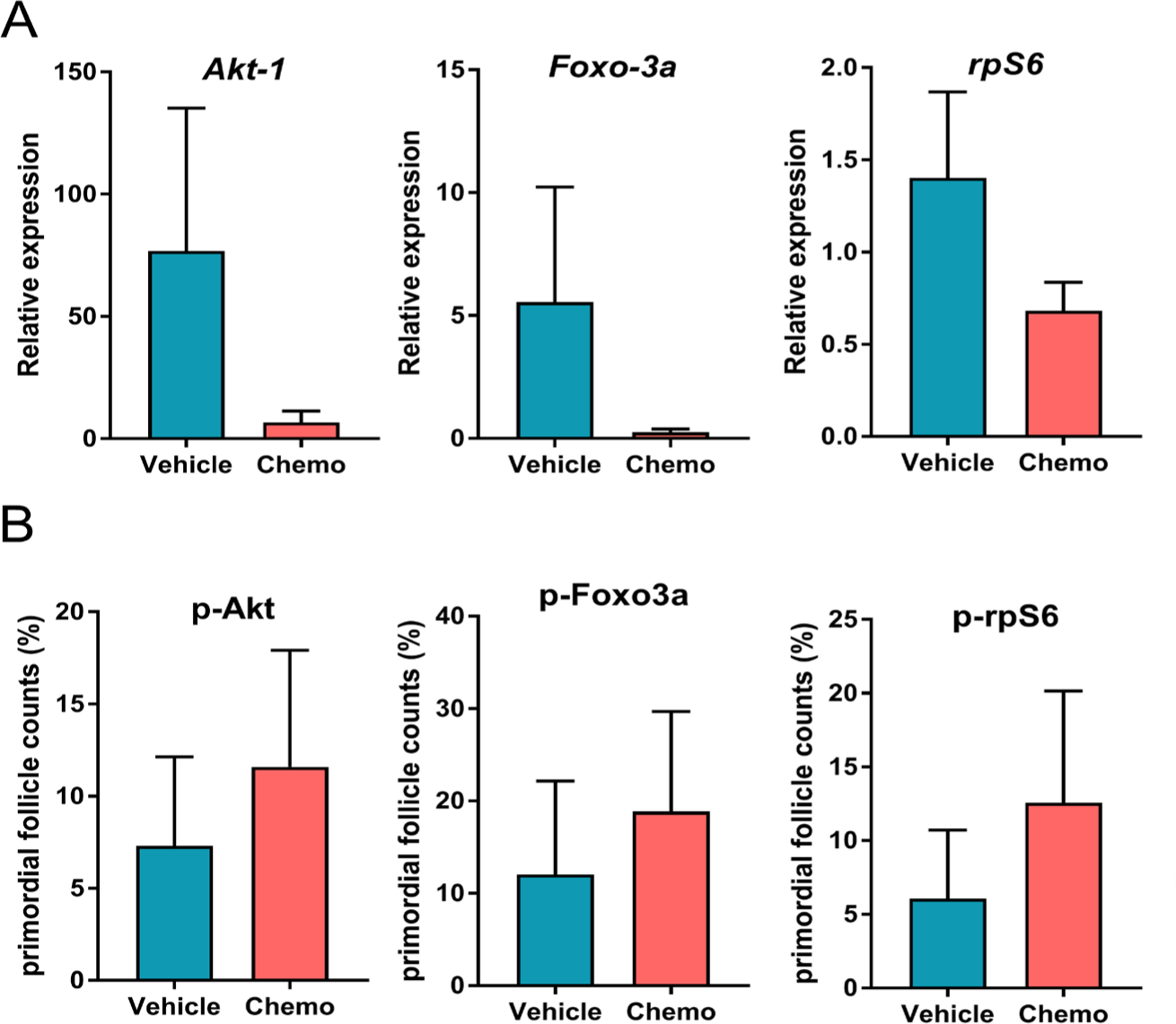
Validation of individual primordial follicle oocyte RNA-seq results by qRT-PCR and immunochemistry. (A) qRT-PCR analysis of PI3K/PTEN/Akt pathway genes shows no significant change in the expression of Akt-1 (n _vehicle_ = 5 and n _chemo_ = 6; p = 0.93), rPS6 (n _vehicle_ = 5 and n _chemo_ = 6; p = 0.32) or Foxo3a (n _vehicle_ = 5 and n _chemo_ = 6; p = 0.12) in the cyclophosphamide-treated group (chemo) as compared to the vehicle. The ‘n’ indicates the number of single primordial follicles analyzed, data are presented as mean of the ΔΔCt ± S.E.M. (B) There is no change in the percentage of primordial follicles stained for (phospho)-Akt (7.29 ± 0.22, n = 8 vs. 11.59 ± 0.8, n = 6; p = 0.52), p-Foxo3a (12.03 ± 1.48, n = 9 vs. 18.85 ± 1.43, n = 8 ; p = 0.63), and p-rpS6 (6.06 ± 0.14, n = 11 vs.12.55 ± 0.94, n = 7; p = 0.4) after cyclophosphamide treatment. The ‘n’ indicates the number of xenografted ovarian cortex tissues, data are presented as mean ± S.E.M.

### Chemotherapy Activates Pro-apoptotic BAD-Bcl2 Action

*Bcl2* (B-cell lymphoma 2) is a known anti-apoptotic gene [23]. It is localized to the outer membrane of the mitochondria, and it plays an important role in promoting cell survival via inhibiting the actions of pro-apoptotic genes such as the *BAD*. By qRT-PCR analysis of laser-captured primordial follicle oocyte we observed a trend towards declining expression of *Bcl2* (n _vehicle_ = 5 and n _chemo_ = 6; p = 0.17), but together with *BAD* (n _vehicle_ = 5 and n _chemo_ = 6; p = 0.93), it did not reach statistical significance in the cyclophosphamide -treated xenografts vs. the vehicles (Figure 5A-B). Under certain stress conditions, BAD is activated by dephosphorylation and dimerizes with BCL2, preventing it from its anti-apoptotic function, and promote apoptosis. We showed a significantly increased colocalization of BAD and BCL2 protein staining in cyclophosphamide -treated samples (n = 6) compared with the vehicle (n = 7; p = 0.004) (Figure 5C, D-J), which may suggest increased protein colocalization and potential interactions. These results collectively indicate that within 12h, chemotherapy exposure results in increased liability to apoptotic death in primordial follicle oocytes, possibly via the BAD-Bcl2 pathway.

**Figure 5.**
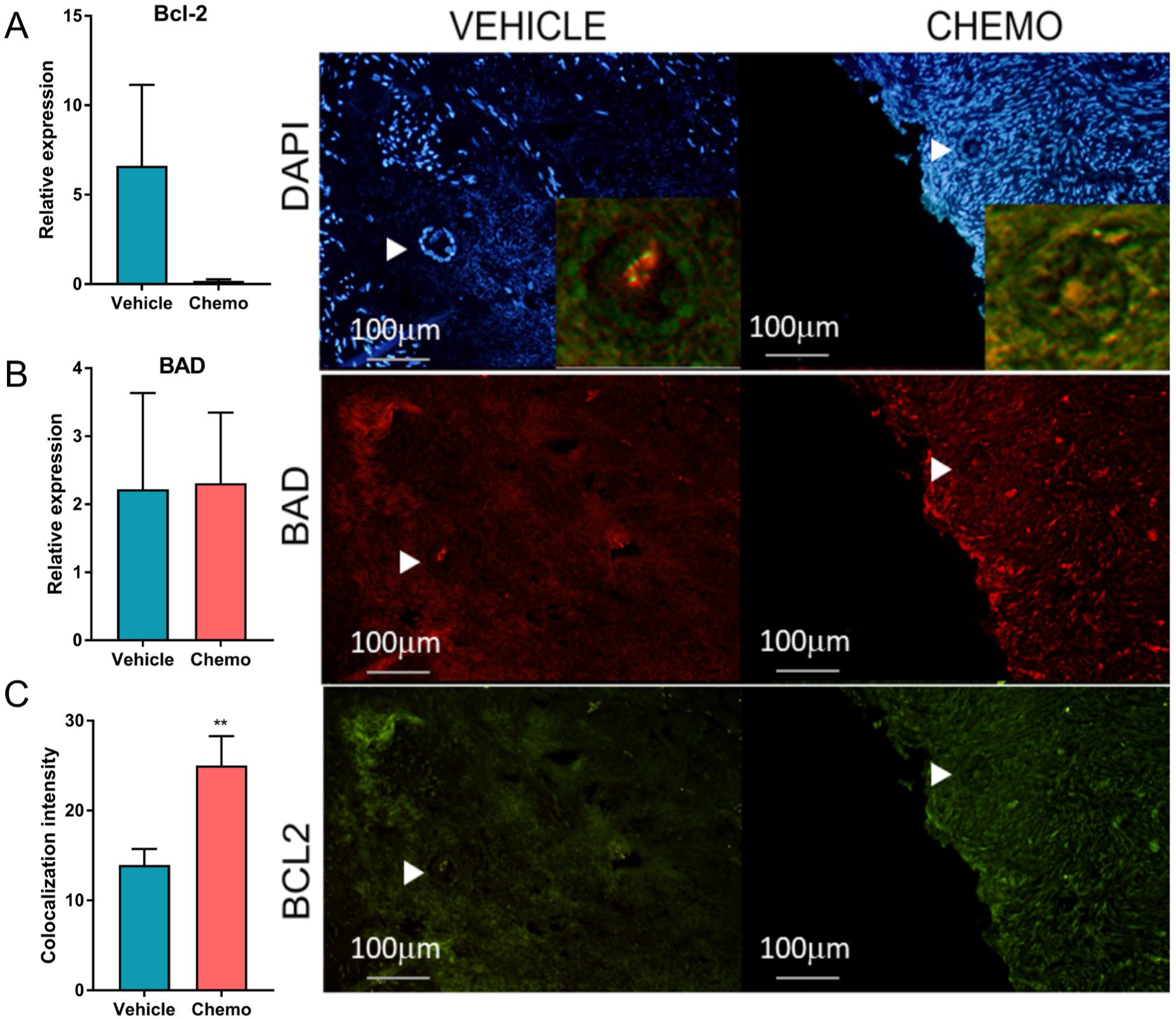
qRT-PCR and immunostaining analysis of BAD and BCL2 12h after cyclophosphamide exposure. qRT-PCR analysis shows no significant change in the expression of both (A) Bcl-2 (n _vehicle_ = 5 and n _chemo_ = 6; p = 0.17) and (B) BAD (n _vehicle_ = 5 and n _chemo_ = 6; p = 0.93) after cyclophosphamide exposure (chemo) compared to the vehicle group. The ‘n’ indicates the number of single primordial follicle oocytes analyzed, data are presented as ΔΔCt mean of the ± S.E.M. (C) By Image J analysis, the BAD-BCL2 colocalization intensity was increased in primordial follicles after cyclophosphamide exposure (n = 6-7; p = 0.004), ‘n’ indicates the number of different xenografted tissue pieces, data are presented as mean ± S.E.M. (D-E) Immunofluorescence images representative of BAD-BCL2 staining in the vehicle- and chemo-treated samples. White arrowheads point to primordial follicles. Inset (J) exemplifies colocalization (yellow color) of BAD and BCL2 in a primordial follicle.

## DISCUSSION

Maintenance of primordial follicle reserve is crucial for the reproductive life span of a female individual. This reserve incurs a massive loss when a patient undergoes chemotherapy treatment, later presenting itself as early menopause and infertility. In this study, we, for the first time, identified the molecular mechanisms of chemotherapy-induced acute damage to ovarian primordial follicles using our unique *in vivo* human ovarian xenograft model and single-cell transcriptomic approaches. Because our earlier work showed that human primordial follicle apoptosis peaks 12 hours post-exposure to chemotherapy [5], and as our focus was to determine the mechanisms of acute follicle loss after cancer treatments, we limited our experiments to 12h time point. We provided multiple lines of evidence converging that cyclophosphamide induces primordial follicle death by prompt activation of apoptotic death pathways, not the PI3K/PTEN/Akt pathway. While we showed that within 12h of treatment, chemotherapy induces DNA DSBs in human oocytes, we found no evidence that it results in increased growth activation of primordial follicles. Likewise, our previous immunohistochemical analyses in rodents and human ovarian xenograft models showed that both cyclophosphamide and doxorubicin induces DSBs and apoptotic oocyte death in primordial follicles [3].

In this unique *in vivo* human ovarian xenograft model, we first showed that chemotherapy exposure did not acutely result in the change of total and non-atretic primary vs. primordial follicle ratio rather, it induced DNA DSBs and AC-3 expression in primordial follicles. This morphological evidence supports that chemotherapy induces primordial follicle apoptotic death rather than growth initiation (Supplementary Figure 2).

The morphological data was then backed by LCM-based individual-oocyte RNA-seq and IPA analysis, which we used to interrogate the key pathways that are activated in primordial follicles in response to chemotherapy. The LCM-based cell isolation method enables us to precisely capture individual primordial follicle oocytes from ovarian tissue where they are identified based on their morphology without the need of unique genetic markers. This initial screening directed us towards pathways related to apoptotic processes rather than those that are involved in follicle growth activation as culprits in acute chemotherapy-induced ovarian reserve loss. Considering that the number of the sequenced primordial oocyte follicles was relatively small, we then confirmed these findings by qRT-PCR and immunohistochemistry in a larger number of samples (n=5-6 and n=8-11, where ‘n’ is the number of single primordial follicle oocytes and number of xenografted ovarian tissues belonging to different donors, respectively).

The transcriptomic analysis with our LCM-based individual-oocyte RNA-sequencing approach indicated suppression of several signaling pathways such as Phospholipase-C (PLC), Ephrin (Eph) Receptor, and IL-8 that are anti-apoptotic and involved with Bad-Bcl2-mediated apoptosis [24][25][26]. Activation of IKBKE induces NF-κB nuclear accumulation and DNA-binding activity by phosphorylation, which then increases the transcription of *BCL2* [27]. Although we did not find a significant decrease in the expression of pro-survival gene BCL-2 and pro-apoptotic BAD, we observed an increased colocalization of BAD with BCL2, which indicates an increased tendency for apoptotic death in human primordial follicle oocytes. We, on the other hand, did not find an increase in the phosphorylation of the key PI3K/PTEN/Akt members in acute response to chemotherapy. These converging findings support the common theme that chemotherapy induces human primordial follicle death via apoptotic pathways rather than massive follicle growth activation and resulting ‘burn-out’ in the acute phase. While our data cannot rule out the activation of the primordial follicles at the later time point, it confirms apoptosis as the acute response of the oocyte to chemotherapy.

In agreement with our findings in humans, other studies in rodents showed that alkylating agents busulfan and cyclophosphamide induce apoptosis and deplete primordial follicle reserve [8,28–31]. Studies in mouse oocytes also showed that cyclophosphamide induces a reduction in mitochondrial transmembrane potential and accumulation of cytochrome-c in the cytosol, leading to activation of the caspase family and apoptosis [32]. A plethora of factors such as DNA damage, energy stress, loss of growth factor signaling and hypoxia can trigger apoptosis by activation of BCl2 pathway proteins [33]. In our previous *in vitro* and xenografting studies, we showed that a topoisomerase inhibitor, cancer drug doxorubicin induces DNA DSBs and apoptosis in primordial follicles [3]. In this study, cyclophosphamide treatment likewise increased DNA DSBs, as shown by increased expression of γH2AX staining in the primordial follicles. Increased DNA damage in primordial follicles is a likely stress signal for the recruitment of the pro-apoptotic BCL2 that causes the activation of the apoptotic cascade in the primordial follicles.

Our individual-oocyte transcriptomic analyses also showed significant downregulation of platelet endothelial cell adhesion molecule-1 (PECAM), and Angiopoietin-1(ANGPT1) (Figure 3D-F). Previous studies have shown that Jurkat cells with a 70% reduction of PECAM-1 expression by siRNA-silencing were significantly more sensitive to chemotherapy-induced apoptosis [34]. Neuronal cell death was caused by the downregulation of Ang-1 (a.k.a. ANGPT1, followed by Akt pathway inhibition [35]. It is also shown that Ang-1 induces phosphorylation of Akt, which is associated with the up-regulation of the apoptosis inhibitor survivin [22]. Inhibition of ANGPT1 in rat follicles increased the levels of AC-3 and decreased Akt phosphorylation [35]. The latter concurs with our finding of increased AC-3 expression after chemo exposure in primordial follicles and brings up the interesting possibility that chemo exposure may indeed result in the suppression of Akt functions, which further favors apoptosis. Interestingly, in addition to PECAM and ANGPT1, IKBKE is also a positive regulator of Akt activation [19][21][22], which we found to have decreased in expression in primordial follicles after chemotherapy.

In summary, our *in vivo* human data show that by suppression of anti-apoptotic mechanisms, gonadotoxic chemotherapy exposure induces a pro-apoptotic state in primordial follicles in the acute phase (Figure 6). Rather than PI3K/PTEN/Akt pathway activation resulting in massive follicle growth and ‘burn-out’, we found contrary evidence that chemotherapy may favor inactivation of the pathway. Nonetheless, our study only focused on the acute effects of chemotherapy. Additional follicle loss may probably occur at later time points as a result of additional mechanisms such as the stromal/microvascular damage and follicle activation. However, given the magnitude of acute damage from DNA damage and apoptotic death, we propose that the research for the development of targeted medical gonadal fertility preservation treatments should focus on preventing DNA damage and/or chemo-induced pro-apoptotic state in primordial follicles.

**Figure 6.**
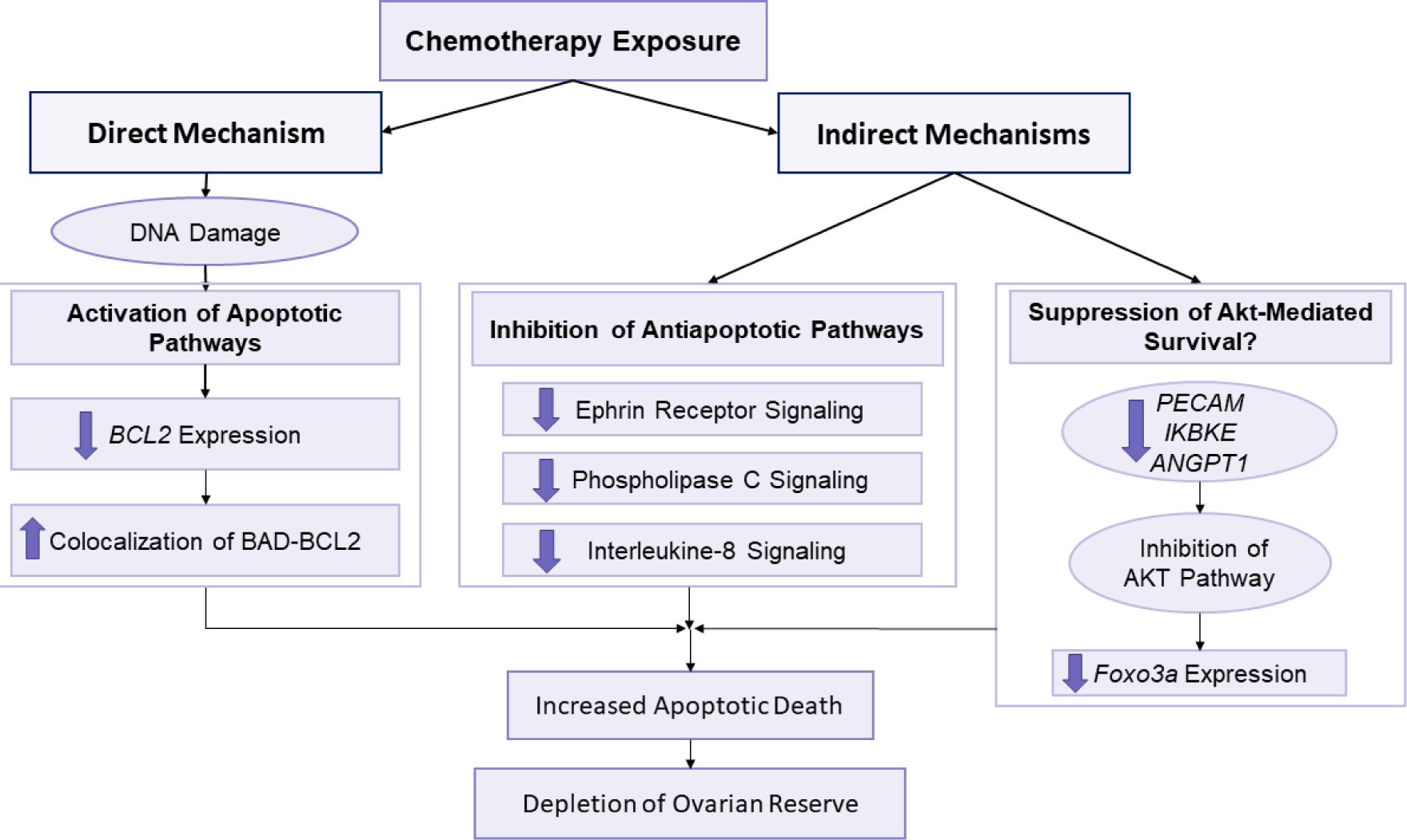
Mechanisms of acute chemotherapy-induced damage to ovarian reserve in human. Chemotherapy exposure induces apoptotic death in primordial follicles by direct and indirect mechanisms, and results in the massive depletion of human ovarian reserve. Directly, gonadotoxic chemotherapy agents such as cyclophosphamide induce double-strand DNA breaks in human primordial follicle oocytes which results in the activation of apoptotic cascades and increased co-localization of BAD-BCL2. Indirectly, same chemotherapy exposure also leads to suppression of anti-apoptotic pathways such as the Ephrin Receptor Signaling, Phospholipase C Signaling, and Interleukin-8 Signaling. Likewise, indirect suppression of pro-Akt pathways such as PECAM-1 (Platelet/Endothelial Cell Adhesion Molecule-1), IKBKE (inhibitor of nuclear factor kappa-B kinase subunit epsilon) and ANGPT1 (Angiopoietin-1) likely leads to the inhibition of the pro-survival functions of the Akt pathway. The sum of the latter two indirect mechanisms is increased sensitivity to apoptotic, gonadotoxic agents. While this scheme explains the mechanisms of acute primordial follicle losses after chemotherapy, there could be other direct and indirect mechanisms responsible for delayed losses, which are not studied here. For more detailed discussion and an expanded figure, please refer to Szymanska *et al*.,2020

## AUTHOR CONTRIBUTIONS

K.O. designed the study and conceived the idea, S.T, B.M, V.T, E.T performed the experiments, S.T and K.S analyzed the results, R.G. assisted with bioinformatic analysis, K.O, S.T., and K.S wrote the manuscript

## ADDITIONAL INFORMATION

### Grant support

Research was supported by R01 HD053112

### Competing Interests Statement

The authors declare no competing interests.

## SUPPLEMENTARY DATA LEGENDS

**Supplementary Figure 1. Detailed representation of altered genes in acute response to chemotherapy exposure in human primordial follicles**. IPA predicted activation of apoptotic processes in the cyclophosphamide-treated samples as there is a decrease in the expression of the genes regulating the phospholipase C, Ephrin and IL-8 signaling. A decrease in the expression of the genes of these anti-apoptotic pathways predicted activation of apoptosis in cyclophosphamides-treated primordial follicles.

**Supplementary Figure 2. Summary of experimental design, methods and results**.

**Supplementary Table 1. qRT-PCR primers for sequencing validation**.

**Supplementary Table 2. Differentially expressed genes between the vehicle- and cyclophosphamide -treated groups**. Fold change ≥ 2, p<0.05.

## Notes

### Competing Interest Statement

The authors have declared no competing interest.

