## Supplementary data for "*In Vivo* Mechanisms of Chemotherapy-Induced Acute Follicle Loss in the Human Ovary: An Individual-Oocyte Transcriptomic Analysis from Human Ovarian Xenografts"

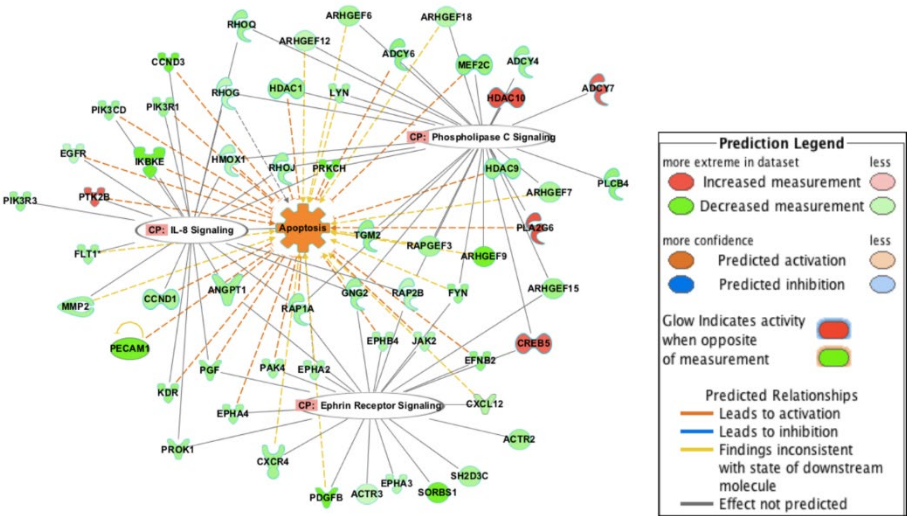


**Supplementary Figure 1. Detailed representation of altered genes in acute response to chemotherapy exposure in human primordial follicles.** IPA predicted activation of apoptotic processes in the cyclophosphamide-treated samples as there is a decrease in the expression of the genes regulating the phospholipase C, Ephrin and IL-8 signaling. A decrease in the expression of the genes of these anti-apoptotic pathways predicted activation of apoptosis in cyclophosphamides-treated primordial follicles.

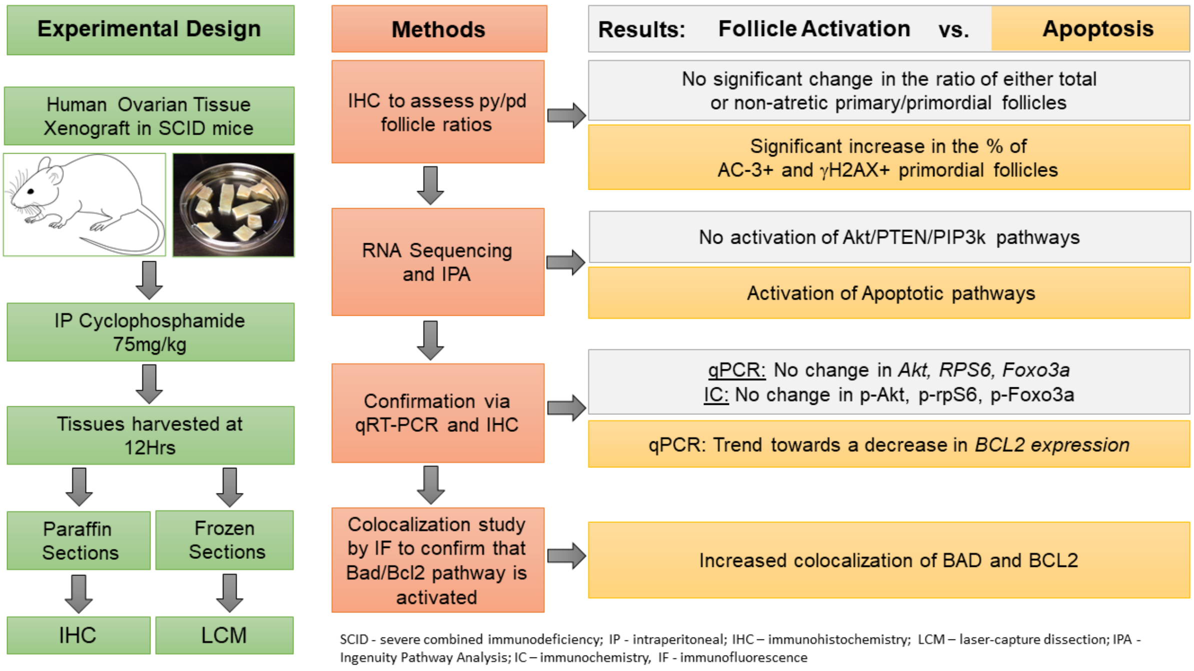


**Supplementary Figure 2. Summary of experimental design, methods and results.**

**Supplementary Table 1. qRT-PCR primers for sequencing validation.**

| Gene | Forward Primer (5' -3') | Reverse primer (5' - 3') | Reference |
|---|---|---|---|
| **βActin** | GGACTTCGAGCAAGAGATGG | AGCACTGTGTTGGCGTACAG | [36] |
| **Akt 1** | ATGAGCGACGTGGCTATTGTGAAG | GAGGCCGTCAGCCACAGTCTGGAT G | [37] |
| **rpS6** | CTGAACATCTCCTTCCCAGCCA | CCTTGTTTGTCGTTCCCACCAC | [38] |
| **FOXO3a** | AAATGAAAGCTCACTCTGGATTCC | TGTGCAATTCCTATGCAATC | [39] |
| **BCL2** | GGTGGGGTCATGTGTGTGG | CGGTTCAGGTACTCAGTCATCC | [40] |
| **BAD** | CCCAGAGTTTGAGCCGAGTG | CCCATCCCTTCGTCGTCCT | [41] |

**Supplementary Table 2. Differentially expressed genes between the vehicle- and cyclophosphamide -treated groups.** Fold change ≥ 2, p<0.05.

| Symbol | log2 Fold Change | P adj |
|---|---|---|
| **UBE3C** | -9.180882449 | 0.040085333 |
| **BTBD7** | -22.25567861 | 2.13E-09 |
| **PRKCH** | -20.51335026 | 3.31E-06 |
| **ARAP2** | -20.4954675 | 2.66E-06 |
| **MSMO1** | -16.0096814 | 0.008788797 |
| **PRDM1** | -10.20430463 | 0.043086696 |
| **MPPED2** | -19.48784036 | 9.01E-05 |
| **ING3** | 35.44168473 | 1.76E-11 |
| **RARB** | -17.30807194 | 0.000306996 |
| **RUNX1T1** | -22.03094937 | 1.74E-07 |
| **EXD2** | -16.9633179 | 0.000103482 |
| **MEF2C** | -10.16269657 | 0.045965997 |
| **COL19A1** | -18.3385071 | 0.001131939 |
| **SLCO1A2** | -19.96074333 | 0.000102151 |
| **ATRN** | -22.03906099 | 3.05E-09 |
| **SORBS1** | -23.7788134 | 2.03E-08 |
| **ERMP1** | -22.52222231 | 1.95E-07 |
| **PDGFB** | -21.26474926 | 1.01E-07 |
| **CACNA1I** | 29.19208883 | 4.34E-08 |
| **HDAC10** | 29.27917335 | 6.38E-08 |
| **SEC23B** | -23.23375918 | 4.17E-10 |
| **POLI** | -21.1819802 | 1.36E-08 |
| **DGKH** | -10.65301456 | 0.017627077 |
| **SLC30A4** | -19.60037782 | 0.000305686 |
| **DECR1** | -20.59045592 | 0.00011553 |
| **ARMC1** | -20.55842663 | 9.18E-08 |
| **CARD8** | -21.88104536 | 2.97E-09 |
| **MAG** | -17.7149294 | 0.001294348 |
| **MPP6** | -21.0782514 | 7.46E-07 |
| **SERPINE1** | 24.35415795 | 4.19E-06 |
| **SUSD1** | -20.4857159 | 0.000253897 |
| **MTMR4** | -22.0869566 | 7.31E-09 |
| **PNPO** | 33.6783901 | 1.61E-15 |
| **ENO3** | 18.94379419 | 0.000608386 |
| **AKAP10** | -10.44955145 | 0.028589419 |
| **CNTNAP1** | -23.15936896 | 2.88E-07 |
| **FRG1** | -18.03911976 | 0.000118365 |
| **CBL** | -9.640487536 | 0.046280656 |
| **VWF** | -12.99874622 | 0.010713752 |

| | | |
|---|---|---|
| **PRDM4** | -23.32442267 | 1.91E-08 |
| **RIC8B** | -20.75051691 | 7.88E-06 |
| **CCND3** | -19.56491465 | 3.04E-06 |
| **LNPEP** | -21.02597365 | 1.09E-06 |
| **UMPS** | -16.24989683 | 0.000160938 |
| **TANC1** | -10.1182097 | 0.030314815 |
| **PDE1A** | -18.27341468 | 0.001166159 |
| **IL18R1** | 20.77140348 | 0.000102144 |
| **TIA1** | -10.24383351 | 0.008582286 |
| **NID1** | -9.92446493 | 0.040085333 |
| **UAP1** | 20.63352781 | 8.93E-06 |
| **PHF19** | -18.0354321 | 0.000937452 |
| **CLU** | -17.79400573 | 1.86E-05 |
| **RNF2** | -10.02982899 | 0.035465596 |
| **GTDC1** | -21.67052739 | 1.92E-06 |
| **ZC3H13** | -9.821040712 | 0.040657763 |
| **PFKFB2** | 17.68006707 | 0.001831132 |
| **FAM124B** | -26.63308197 | 1.79E-06 |
| **NCOA3** | -24.18374213 | 4.66E-11 |
| **CSE1L** | -10.13920406 | 0.049144593 |
| **EFNB2** | -10.78362224 | 0.040657763 |
| **CHCHD5** | -18.77963693 | 7.99E-06 |
| **CD93** | -10.81062304 | 0.048661808 |
| **CASD1** | -20.24726369 | 1.54E-06 |
| **ELL3** | -18.49467853 | 0.000213266 |
| **OLFM1** | 20.96703376 | 0.000108969 |
| **LAMA5** | -11.28066061 | 0.010797172 |
| **ARHGEF9** | -21.31265053 | 1.09E-08 |
| **NES** | -24.06359042 | 4.19E-09 |
| **PEMT** | 17.21280874 | 0.000366853 |
| **RFC3** | -15.64003897 | 0.003353584 |
| **ADAM30** | 15.06788772 | 0.030373776 |
| **RSAD2** | -18.80106 | 0.000116153 |
| **OSTF1** | -18.01732653 | 0.000134545 |
| **ADAM19** | -21.55762564 | 4.84E-07 |
| **ZC3H10** | -19.67488902 | 4.70E-05 |
| **MAP7** | -17.2512527 | 0.002866974 |
| **ATAT1** | -19.38991174 | 2.26E-06 |
| **SEMA6D** | -22.8897922 | 6.84E-08 |
| **SPTBN5** | -20.11399628 | 6.96E-06 |
| **GCOM1** | 20.07923462 | 2.34E-05 |
| **ENPEP** | -20.94283703 | 1.04E-05 |
| **N4BP2L1** | -17.88019831 | 4.54E-05 |

| | | |
|---|---|---|
| **GREB1L** | -18.29158073 | 0.000669577 |
| **HSPG2** | -7.931575904 | 0.019330356 |
| **MOB3C** | -10.14987715 | 0.034658932 |
| **CTTNBP2NL** | -20.19057034 | 1.17E-06 |
| **RGS5** | -30.25289435 | 1.93E-09 |
| **HMCN1** | -9.689614575 | 0.042891939 |
| **GOLPH3L** | 20.46109531 | 1.33E-06 |
| **LYST** | -9.828521564 | 0.046280656 |
| **CEP170** | -21.78388943 | 2.77E-09 |
| **ATG12** | -9.792294539 | 0.042891939 |
| **ARHGAP26** | -20.32256738 | 6.66E-07 |
| **ZNF300** | -23.72328262 | 9.83E-11 |
| **MUT** | 20.18831251 | 1.93E-05 |
| **VWDE** | -22.09349921 | 4.71E-08 |
| **NRBF2** | 21.13521377 | 5.76E-06 |
| **IGSF10** | -17.58703584 | 0.000411629 |
| **SPARCL1** | -22.23919408 | 5.51E-08 |
| **MR1** | 18.49995251 | 0.00011994 |
| **RGPD3** | -16.08061108 | 0.007400237 |
| **AFAP1L1** | -30.17985081 | 4.17E-10 |
| **MPZL3** | -15.9565452 | 0.006328824 |
| **BBS5** | 17.00645382 | 0.002974405 |
| **ATP1A1** | -10.02617289 | 0.034931684 |
| **EMCN** | -30.89778162 | 3.09E-10 |
| **AL353743.1** | -18.6023219 | 0.000359661 |
| **ARMC3** | -16.67429762 | 0.0031464 |
| **C10orf10** | -23.44177534 | 2.92E-10 |
| **ZNF143** | -20.35430683 | 2.20E-06 |
| **MCM7** | -23.85132366 | 9.80E-09 |
| **TERF2IP** | -21.134117 | 3.40E-07 |
| **TRANK1** | -24.35123465 | 2.02E-10 |
| **REEP4** | 34.4432497 | 3.81E-11 |
| **TSPAN5** | -21.2573024 | 4.18E-07 |
| **SLC35G2** | -30.39454241 | 1.36E-10 |
| **LDB2** | -30.66872459 | 1.33E-08 |
| **MAP3K2** | -10.01730122 | 0.047922213 |
| **COMMD5** | -21.8663568 | 2.67E-06 |
| **ZNF16** | -21.07545007 | 1.62E-07 |
| **KBTBD2** | -10.26573269 | 0.033774936 |
| **KCND3** | -17.8810868 | 0.001166159 |
| **SLC19A1** | -12.28347574 | 0.001308306 |
| **ADCY6** | -10.55009928 | 0.010073599 |
| **CMKLR1** | -17.78361551 | 0.000289413 |

| | | |
|---|---|---|
| **PAAF1** | -21.34019035 | 4.84E-07 |
| **CCDC57** | -19.4782915 | 2.52E-07 |
| **SGF29** | 21.19938757 | 9.56E-09 |
| **RIMKLA** | -21.56671249 | 3.07E-06 |
| **GPC5** | -15.77015927 | 0.010599268 |
| **DNHD1** | -21.12993266 | 2.36E-08 |
| **ANKRD18A** | 30.72388602 | 8.59E-09 |
| **ZDHHC20** | -22.05966297 | 2.24E-06 |
| **RNF135** | -16.92388324 | 0.004277318 |
| **DEXI** | -17.56646109 | 0.001831132 |
| **ADGRG3** | 16.7874542 | 0.002302511 |
| **ZFP1** | -20.52784498 | 5.34E-06 |
| **KIAA0825** | -18.93213716 | 0.000305518 |
| **GPRIN3** | -19.13502957 | 1.24E-05 |
| **PRKG1** | -32.83887845 | 1.29E-10 |
| **LAMP1** | -21.89685688 | 2.40E-09 |
| **ANKRD37** | 17.68434945 | 5.96E-05 |
| **TNFRSF4** | -19.81579227 | 0.000389422 |
| **ZNF559** | -9.720558653 | 0.046280656 |
| **NDOR1** | -17.74098876 | 0.000181783 |
| **ZNF548** | -18.70994871 | 1.15E-06 |
| **S100A4** | -21.15527224 | 0.0002424 |
| **DACT3** | -18.60073437 | 0.000253897 |
| **DCHS2** | 22.13441408 | 3.93E-06 |
| **ENTPD7** | -21.02901966 | 9.22E-07 |
| **FAM19A2** | -16.36225354 | 0.003249114 |
| **COL15A1** | -11.79653071 | 0.016588591 |
| **ABHD16A** | 21.2288493 | 0.000141662 |
| **GCNT6** | 26.26270332 | 2.55E-06 |
| **CRYZL1** | -10.19429058 | 0.031644741 |
| **SEC14L1P1** | 17.59586167 | 0.000366705 |
| **PLEKHM1P1** | -19.7226533 | 2.26E-06 |
| **LINC00680** | -19.75272435 | 0.000123171 |
| **RPL17-C18orf32** | -18.07827757 | 0.003007074 |
| **AC068491.1** | -29.63228622 | 3.54E-08 |
| **OR52I2** | 29.73227328 | 9.17E-09 |
| **AC092641.1** | 29.88828572 | 2.08E-08 |
| **SATB1-AS1** | -19.50370225 | 3.24E-05 |
| **SNORA71B** | -16.83514163 | 0.007730674 |
| **AL035448.1** | -21.82599564 | 3.30E-06 |
| **MIR600HG** | -19.91330603 | 2.41E-05 |
| **PRMT5-AS1** | -17.38168289 | 0.000267746 |
| **AP001992.1** | -18.65959609 | 0.000102723 |

| | | |
|---|---|---|
| **ACTG1P13** | 29.25362388 | 2.27E-08 |
| **MCPH1-AS1** | 32.28278911 | 5.39E-11 |
| **AC026904.3** | -26.13288878 | 2.81E-06 |
| **AC091053.1** | -21.32092916 | 4.03E-07 |
| **CTSO** | -18.84393824 | 9.01E-05 |
| **TAS2R31** | -15.71595565 | 0.015553005 |
| **TMC3-AS1** | -16.03332764 | 0.003047062 |
| **TYRO3P** | 18.18361836 | 0.000461693 |
| **AC009093.1** | 10.27179944 | 0.047922213 |
| **AC127459.1** | -20.0885839 | 1.09E-06 |
| **PECAM1** | -25.19837655 | 2.13E-09 |
| **PAN3-AS1** | -17.41081947 | 0.001976919 |
| **IKBKE** | -16.9791079 | 0.001656132 |
| **AP000654.1** | 26.42316847 | 1.55E-06 |
| **ZNF2** | -34.49614293 | 5.32E-11 |
| **PADI6** | 25.65424808 | 4.55E-06 |
| **HIST1H4E** | -19.52678387 | 1.12E-05 |
| **BACE1-AS** | -16.08386962 | 0.000768391 |
| **AP000866.6** | -21.67792921 | 2.85E-05 |
| **AC008536.3** | -21.52395032 | 2.67E-06 |
| **AL590434.1** | -20.66578411 | 0.000119772 |
